# PDGF-BB overexpression in p53 null oligodendrocyte progenitors increases H3K27me3 and induces transcriptional changes which favor proliferation

**DOI:** 10.1101/2024.05.14.594214

**Authors:** Dennis Huang, Angeliki Mela, Natarajan V. Bhanu, Benjamin A. Garcia, Peter Canoll, Patrizia Casaccia

## Abstract

Proneural gliomas are brain tumors characterized by enrichment of oligodendrocyte progenitor cell (OPC) transcripts and genetic alterations. In this study we sought to identify transcriptional and epigenetic differences between OPCs with *Trp53* deletion and PDGF-BB overexpression (BB-p53n), which form tumors when transplanted in mouse brains, and those carrying only p53 deletion (p53n), which do not. We used unbiased histone proteomics and RNA-seq analysis on these two genetically modified OPC populations and detected higher levels of H3K27me3 in BB-p53n compared to p53n OPCs. The BB-p53n OPC were characterized by higher levels of transcripts related to proliferation and lower levels of those related to differentiation. Pharmacological inhibition of histone H3K27 trimethylation in BB-p53n OPC reduced cell cycle transcripts and increased the expression of differentiation markers. These data suggest that PDGF-BB overexpression in p53 null OPC results in histone post-translational modifications and consequent transcriptional changes favoring proliferation while halting differentiation, thereby promoting the early stages of transformation.

## Introduction

Gliomas are the most common malignant tumor in the central nervous system (CNS). They are characterized by rapid disease course and a marked cellular^1,2,3,4,5^, transcriptional and epigenetic^6,7^ heterogeneity. This described complexity likely contributes to the difficulty of advancing treatment options and developing viable drug options^8,9,10^, rendering these tumors incurable.

In an attempt to decipher glioma heterogeneity, previous studies on gene expression profiling^11,12^ identified the presence of distinctive transcriptional signatures, defining four specific glioma subtypes, on the basis of the abundance of cell-specific patterns of gene expression^13^. One such subtype is the proneural glioma, characterized by the transcriptional signature of oligodendrocyte progenitor cells (OPCs) which are the precursors to myelinating oligodendrocytes and represent the most abundant population of proliferative cells in the adult brain^14,15,16^. Integrated genomic analyses^13,17,18^ further revealed the prevalence of specific genetic mutations in distinct subtypes, with the proneural subtype being characterized by loss of tumor suppressor P53 and amplification of platelet derived growth factor (PDGF) signaling^13^. Several studies of gliomagenesis, in animal models with distinct genetic alterations in diverse brain cell types^19,20^ suggested that OPCs act as cell of origin for proneural gliomas^21,22,23,24,25^. However, the early steps leading from OPCs to tumor formation remain elusive.

In this study, we sought to define the epigenetic and transcriptional changes induced in normal OPCs by two genetic mutations frequently detected in proneural gliomas: p53 deletion and increased PDGF signaling. While PDGF-AA is the primary mitogen for OPCs during brain development^26,27,28^ and PDGF-BB has been related to neovascularization^29,30^ both mitogen isoforms have been reported to be secreted by glioma cell lines^31,32,33^, with PDGF-BB frequently detected in aggressive high-grade glioma^34,35,36^. Here, we use unbiased histone proteomics and transcriptomic analysis to assess the intrinsic differences between OPCs carrying p53 deletion, which do not form tumors when injected in mice, and OPCs with p53 deletion and overexpression of PDGF-BB, which form tumors. Our results identify differences in histone H3K27 trimethylation as a regulator of OPC differentiation and proliferation during early stages of gliomagenesis.

## Results

### Cultured OPCs lacking p53 expression and over-expressing PDGF-BB are characterized by increased proliferation and form tumors when injected in mice

The model chosen relies on the use of retroviral vectors and of a *Trp53 flox/flox* transgenic mouse line^37^ (JAX stock #008462). OPCs were isolated from the brains of these transgenic mice and cultured in chemically defined medium prior to the infection with a retroviral vector expressing only the recombinase cre or a bicistronic vector allowing also for PDGF-BB expression^24,38^. This generated OPCs with p53 deletion alone (p53n) or also expressing PDGF-BB (BB-p53n). To assess differences in the proliferative rate between p53n and BB-p53n OPCs, cells were processed for immunocytochemistry, using antibodies specific for OLIG2, as marker of oligodendrocyte lineage cells, and for Ki67, as a marker for proliferation^39,40^. While all the cultured OPC stained for OLIG2, a greater percentage of OLIG2+ were also strongly immunoreactive for Ki67 in BB-p53n OPC cultures (x= 79% +/-6.81% in 30 total cells counted), than in the p53n OPC cultures (x= 28% +/-8.89% in 30 total cell counted) (Fig1B, C). These cells were then subcortically injected into recipient mice, and only those receiving the BB-p53n OPCs formed tumors by 35 days post injection (35dpi), while those injected with p53n OPCs did not reveal any sign of tumor formation, even after 4 months of observations. Accordingly, survival experiments revealed that mice injected with BB-p53n OPCs succumbed to tumor morbidity by 60 dpi, while those injected with p53n OPCs survived past 120 dpi (Fig1D). Histopathology of BB-p53n injected brain tissue at end stage, revealed tumors with features of high-grade glioma, including dense neoplastic growth with pseudo-palisading necrosis and vascular proliferation (Fig 1E). Together, these experiments suggest that PDGF is driving tumor formation via its mitogenic effects of OPCs and led us to compare differences in the transcriptome and epigenetic histone marks between the more proliferative/tumorigenic p53n-BB OPCs and the less proliferative/non-tumorigenic p53nOPCs.

**Figure 1.**
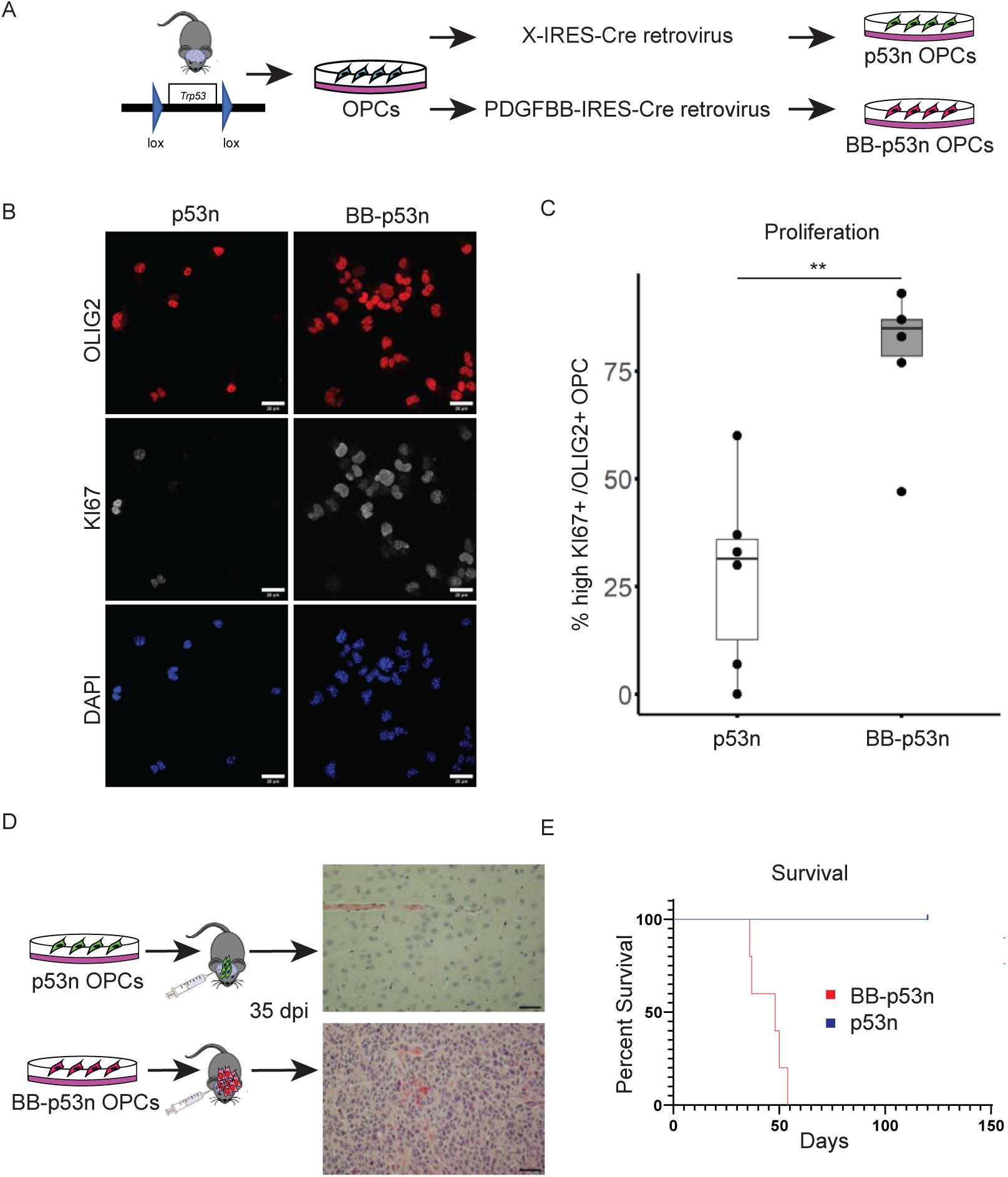
Cultured p53 null OPC over-expressing PDGF-BB are highly proliferative and form aggressive tumors when injected in mice. (A) Schematics of mutant OPCs generation. OPCs cultured from *Trp53 floxed* transgenic mouse were infected either with Cre alone retrovirus (p53n) or with a bicistronic virus expressing both Cre and PDGF-BB (BB-p53n). (B) Confocal images of cultured OPC lacking p53 alone (p53n) or together with PDGF-BB expression (BB-p53n), stained for OLIG2 (red), Ki67 (gray), and DAPI as nuclear counterstain (blue). Scale bar = 20 μm. (C) Boxplot representing the percentage of proliferating (Ki67+/OLIG2+) cells described in 1B. Margins represent the lower and higher quartiles and ends minimum and maximum values of the data obtained from 6 independent determinations (n=30 cells counted for each experiment performed in triplicate in two independent biological samples). High Ki67 determined by nuclear mean fluorescent value above the median value of all samples. Significance two-tailed Student t-test (*p* value = *0.001*). BB-p53n OPC mean percent Ki67+ 79% +/-6.81% SEM. p53n OPC mean percent Ki67+28% +/-8.89% SEM. (D) Schematic and H&E images of p53n and BB-p53n OPCs injected in wild type mice. Only BB-p53null OPC induced histological signs of glioma, as assessed by histopathology after 35 days post injection (dpi). Histology image viewed at 200X magnification, scale bar = 100 μm. (E) Survival plot of mice injected with the OPCs described in 1D (n = 5 mice per group). The blue curve represents p53n mice and the curve represents the BB-p53n mice. Mantel-Cox test used to calculate p-value (*p-* value = *0.0018*)

### The transcriptome of BB-p53null compared to p53null OPCs reveals increased cell cycle transcripts and decreased markers of OPC differentiation

To further investigate whether PDGF-BB overexpression modified the transcriptome of p53n OPCs, we performed RNA-seq. Differential expression analysis between BB-p53n and p53n OPCs revealed over 4000 significantly differentially expressed genes (Supp. Table 1). Among the transcripts with higher expression in the BB-p53n OPC, we detected genes encoding for positive cell cycle regulation (e.g. *Cdc6, Cdc45, Ccne2*) (Fig2A). GSEA analysis^41^ of the upregulated genes confirmed a significant enrichment in gene sets related to proliferation, including those regulating the G1-S and the G2-M transition^42^ (Fig2C), as well as ontologies related to DNA replication and cell division^43,44^ (Fig 2B). Among the downregulated transcripts, we detected several genes related to OPC differentiation (e.g. *Cnp, Myrf. Mbp*), fatty acid metabolism and central nervous system myelination, a major feature of mature oligodendrocytes (Fig 2A,B,D). In addition, BB-p53n OPCs showed a transcriptional signature characterized by altered expression of enzymes responsible for post-translational modifications of histone tails compared to p53n OPC (Fig2E). For instance, transcripts for the enzymes responsible for repressive methylation of histone H3K27 (e.g. *Ezh2, Eed, Suz12*) were upregulated, while those responsible for the removal of this mark (e.g. *Kdm6a, Kdm6b)*, were significantly downregulated. These data support the interpretation that overexpression of PDGF-BB in p53n OPCs enhances their proliferative capabilities, while inhibiting differentiation pathways.

**FIGURE 2:**
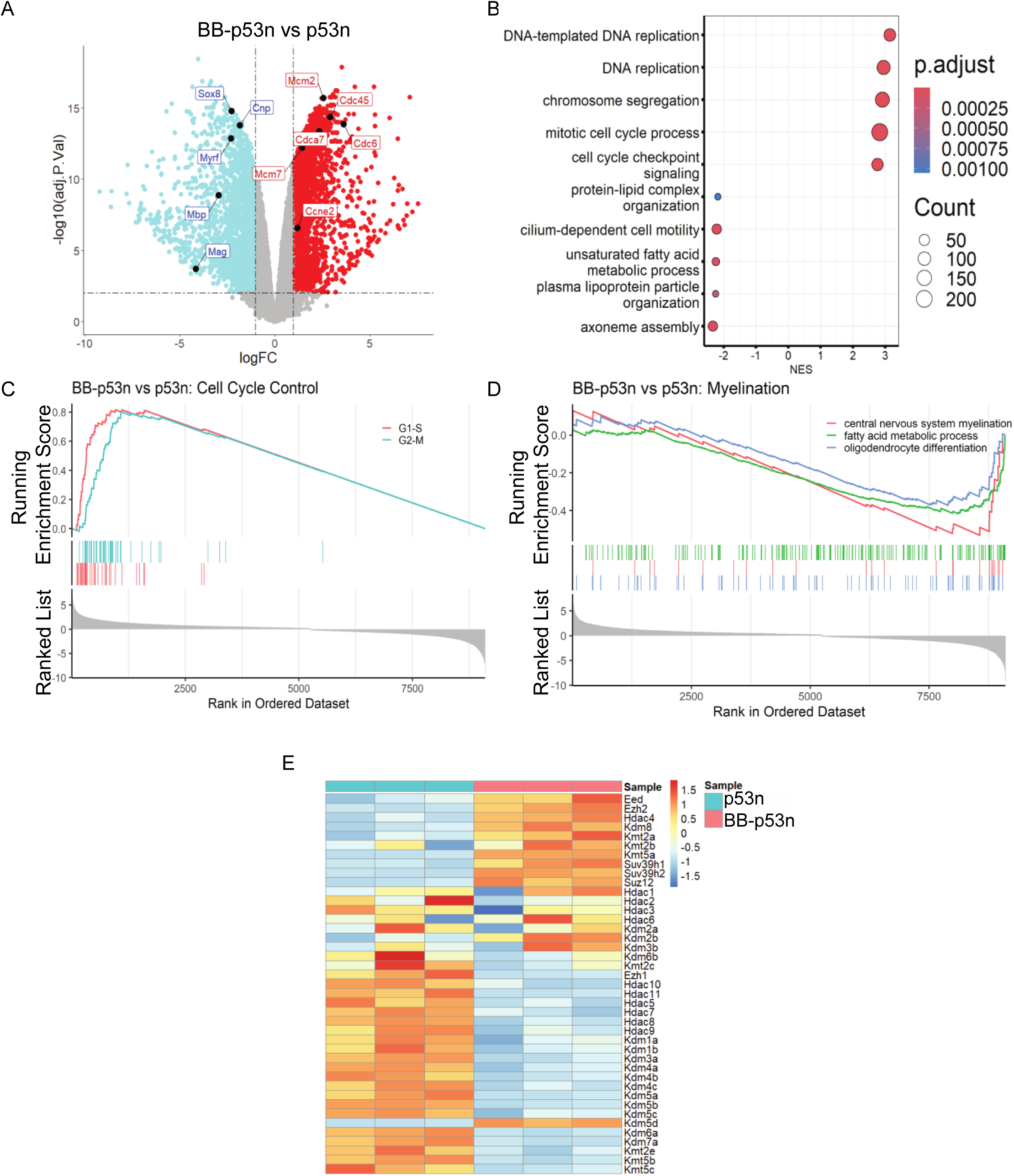
Altered transcriptome in BB-p53n OPCs compared to p53n reveals increased cell cycle transcripts, decreased markers of OPC differentiation and differential expression of histone modifying enzymes. (A) Volcano plot of differentially expressed genes (adjusted p-value cutoff of 0.05 and log_2_ fold change of 1) between BB-p53n and p53n OPCs. Of the 2149 upregulated genes (red), the ones positively regulating cell cycle are indicated in text boxes (red text). Of the 2312 downregulated genes (blue), those related to differentiation are indicated (blue text). (B) Dotplot of GSEA data visualizing multiple GO gene sets, obtained by running the differential expression data against the full GO database. Red to blue color gradient represents adjusted *p*-value. Size of dots represents the number of genes matching between gene set and input gene list. Normalized enrichment score (NES) is plotted on the x-axis. See supplemental materials for full enrichment data table. (C) GSEA of the differential expression data shown in Fig 3A identifying positive regulation of G1-S and G2-M transition. Curated gene sets obtained from Tirosh et al. 2016. Adjusted p-values < 1e-10. (D) GSEA of differential expression data shown in Fig 3A highlighting decreased expression of genes related to OPC differentiation from Gene Ontology database (GO:0048709) adjusted *p*-values = 0.07, central nervous system myelination (GO:0022010) adjusted *p*-value = 0.06, and fatty acid metabolic process (GO:0006631) adjusted *p-*value < 0.001. (E)Heatmap of gene expression data filtered for histone lysine modifying enzymes. Red indicates relatively higher gene expression and blue indicates relatively lower represented by z-scores. Z-score calculated with tpm normalized gene counts.

### BB-p53n OPCs are characterized by higher levels of H3K27me3 than p53n OPCs

The detection of altered transcript levels of histone modifying enzymes led us to hypothesize the presence of higher levels of repressive histone post-translational modifications (e.g. H3K27me3), which could be responsible for the transcriptional changes. We therefore performed unbiased histone proteomics on histones extracted from cultured p53n and BB-p53n OPCs, using mass spectrometry^45,46,47^. Although the differences in histone modifications were mild and did not meet statistical significance, the unbiased histone proteomic analysis suggested increased levels of H3K27me3 in BB-p53n OPC compared to p53n OPCs (Fig 3A). The levels of H3K27me3 were therefore assessed in the two cell populations, using immunocytochemistry (ICC) and western blotting of histone extracts and antibodies highly specific for the histone marks. The nuclear mean fluorescent intensity of H3K27me3 immunoreactivity (Fig. 3B) was quantified, using ImageJ, and revealed higher values in BB-p53n OPCs than p53n OPCs (Fig 3C). Western blot analysis of histones extracts from these two cell population, were also probed with antibodies specific for H3K27me3 and for total H3 and the immunoreactive bands quantified and normalized. Also in this case, BB-p53n OPCs were characterized by significantly higher H3K27me3 levels than p53n OPCs (Fig3D,E). These data suggested that histone modifications observed in glioma cells may occur early in the process of transformation and we therefore hypothesized that the increased levels of H3K27me3 may play a significant role in the transcriptional changes induced by PDGF-BB overexpression in p53 null OPCs.

**FIGURE 3:**
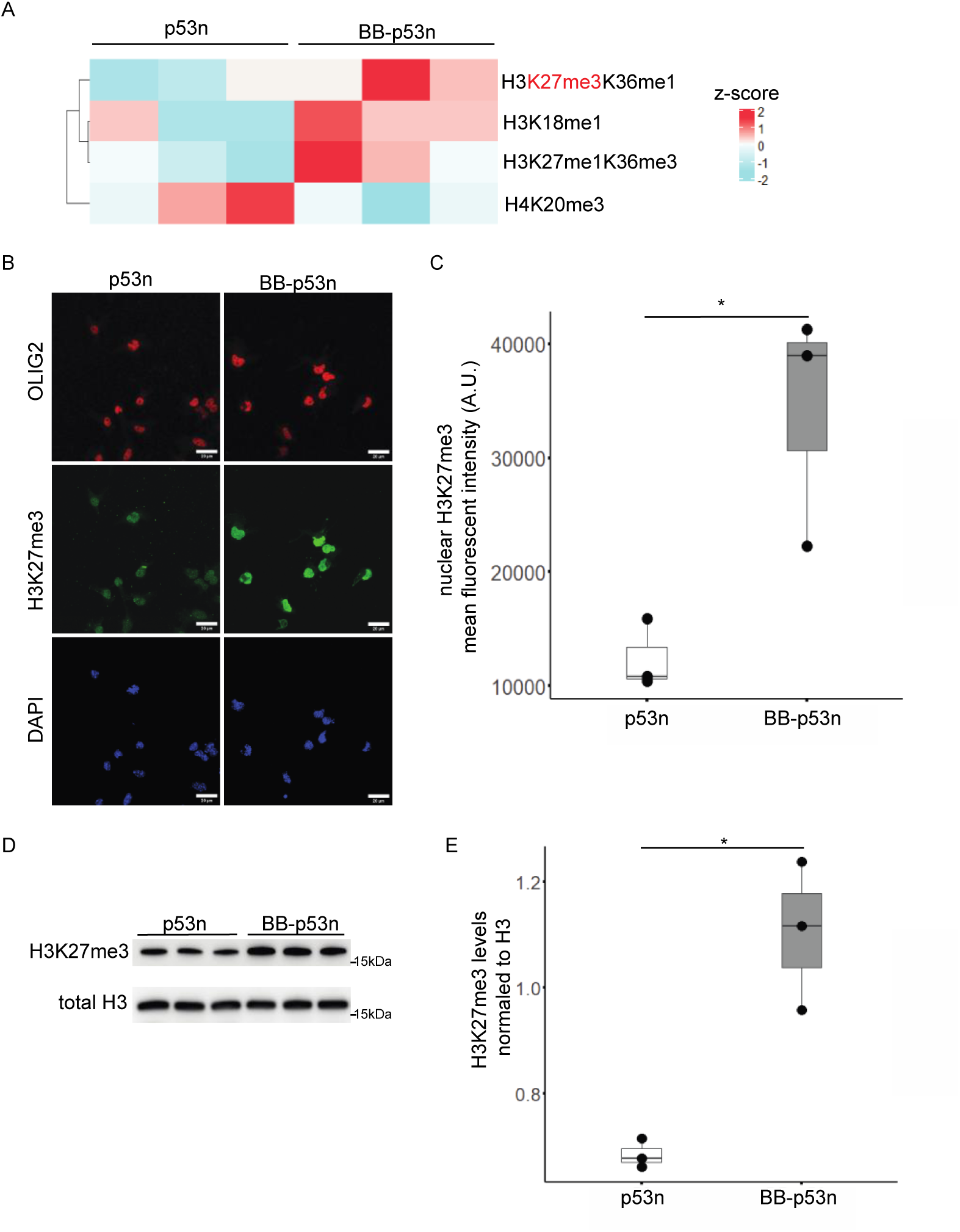
BB-p53n OPCs are characterized by higher levels of H3K27me3 than p53n OPCs. (A)Heatmap of histone modification ratios generated from acid extracted histones and mass spectroscopy. Ratios were converted into z-scores between p53n and BB-p53n samples (see supplemental). Rows organized into unique combinations of lysine modifications in H3 histone tails. Red indicates relatively higher and blue relatively lower levels in OPC of the indicated genotype. (B) Confocal images of cultured p53n and BB-p53n OPCs stained for OLIG2 (red), H3K27me3 (green), and DAPI as nuclear counterstain (blue). Scale bar = 20 μm. (C) Boxplots representing the quantification of nuclear H3K27me3 in Olig2 positive in vitro OPCs visualized in Fig 2B. Margins represent the lower and higher quartiles and ends minimum and maximum values of the data from 3 biological replicates and 30 cells each replicate counted for analysis. p53n mean nuclear fluorescence H3K27me3 = 12377 +/-1758 SEM. BB-p53n mean nuclear fluorescence H3K27me3 = 34182 +/-6000 SEM. Significance calculated by one-tailed student t-test. *p-*value = 0.029. (D) Western blot analysis of acid extracted histones from cultured p53n and BB-p53n OPCs. Histones probed for H3K27me3 and total H3. (E) Boxplot represents the ratio of H3K27me3 over total H3 calculated by dividing the intensity of the H3K27me3 bands by the intensity of total H3 bands from panel 1D. Margins represent the lower and higher quartiles and ends minimum and maximum values of the data. p53n mean normalized H3K27me3 = 0.68 +/-0.02 SEM. BB-p53n mean normalized H3K27me3 = 1.10 +/-0.08 SEM. Significance calculated by two-tailed student t-test (*p-*value = 0.016).

### Pharmacological inhibition of EZH2 in BB-p53n OPC reduces proliferation while increasing differentiation

Considering the significant increase of H3K27me3 in PDGF-BB in p53n OPCs, we reasoned that inhibition of the enzyme responsible for trimethylation of H3K27 could reverse the differential gene expression between these cells, which form tumors when injected in mice, and p53n OPC, which do not form tumors. Since the enzyme EZH2 is responsible for the deposition of the H3K27me3 mark, we treated BB-p53n OPCs with the pharmacological inhibitor of EZH2, Tazemetostat (EZH2i) for 48 hours and validated the effectiveness of the treatment by detecting lower H3K27me3 levels in the EZH2i treated cultures compared to BB-p53n cultures treated with DMSO, as vehicle control (Fig4. A-B). Treated cells (BB-p53n-EZH2i) and their controls (BB-p53n DMSO) were also processed for bulk RNA sequencing. Full transcriptome analysis revealed that the majority of genes in BB-p53n-EZH2i OPCs were not reversed by EZH2 inhibition compared to p53n OPCs (Supp Fig 1A). However, the initial analysis showed a fraction of the gene expression being reversed to show similarity to the expression profile of p53n OPCs. Visualizing those select genes as a heatmap, revealed the presence of three clusters, based on the response to EZH2 inhibition (Fig. 4C). Clusters 1 and 2 included genes regulating cell cycle, mitosis and chromosome segregation, which were expressed at higher levels in the BB-p53OPC compared to p53n OPC, and whose expression was lowered by treatment with the EZH2i (Fig. 4C-D). Cluster 3 included genes related to lipid metabolism, which were expressed at lower levels in the BB-p53n OPCs compared to p53n, were upregulated by treatment with the EZH2i (Fig. 4C-D). As validation of these results we selected transcripts from those clusters for validation by rt-PCR, such as *Ccne2*, the gene encoding for cyclin E2 and *Atad2*, encoding an ATPase known to induce the expression of other cell cycle regulators^48^, and positively regulate the G1-S transition. The results confirmed the higher transcript levels in BB-p53n OPCs compared to p53n and a statistically significant reduction after 48 hours of treatment with the EZH2 inhibitor (Fig4E). In addition, even though the GSEA analysis of OPC differentiation gene set showed insignificant enrichment (data not shown), the transcript levels of genes involved in OPC differentiation such as: *Prmt5*, an arginine methyl transferase important for OPC differentiation^49^ and *Erbb2*, a tyrosine kinase involved in OPC differentiation^50^, showed a trend towards increased expression after 48 hours treatment with EZH2i (Supp Fig 1B). These data suggest that EZH2 inhibition of BBp53n OPCs significantly decreased the expression of genes regulating proliferation, while only partially affected the increase of genes associated with differentiation.

**FIGURE 4:**
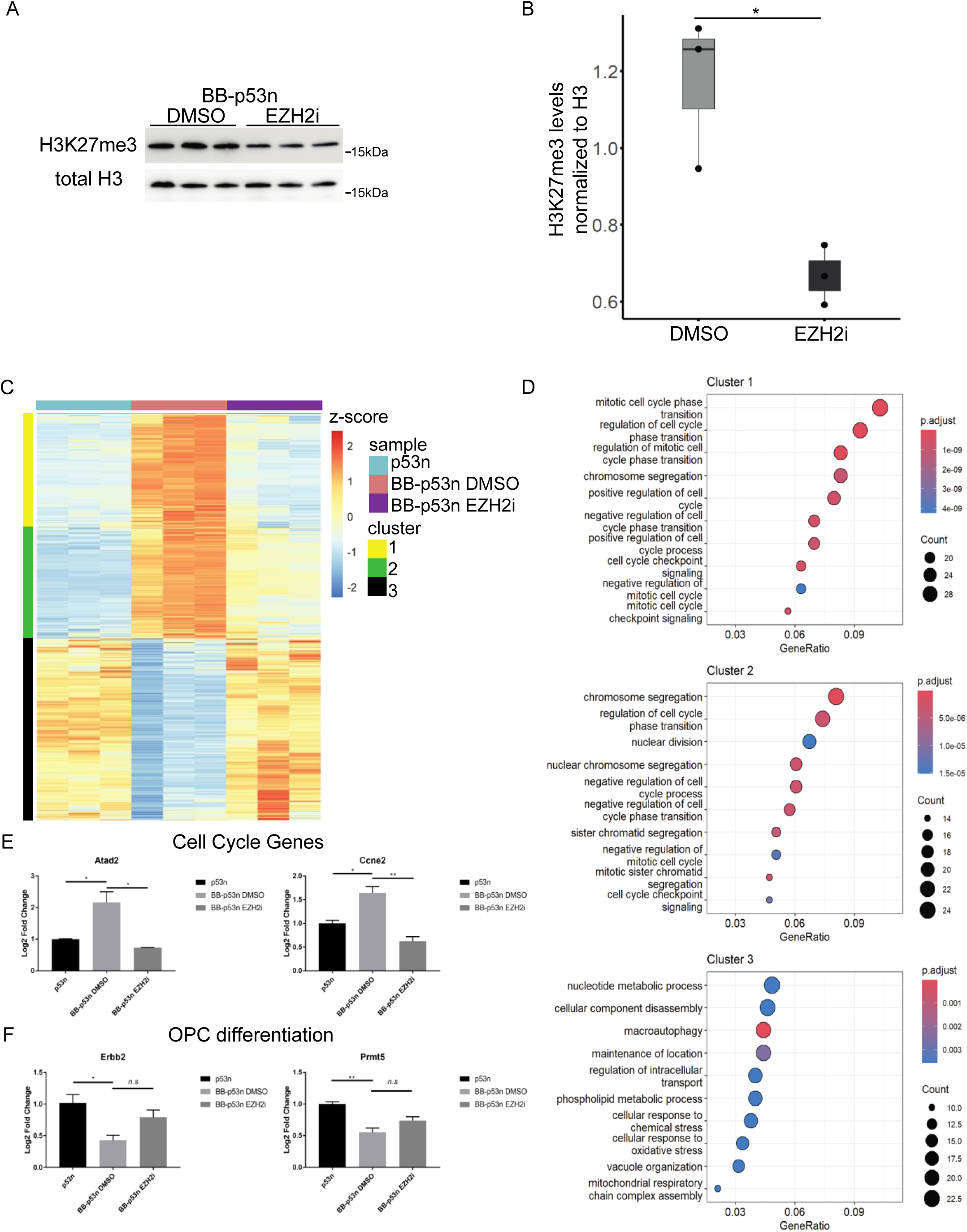
Reduction of H3K27me3 in BB-p53n OPC using pharmacological inhibitors of EZH2 reduced the levels of cell cycle-related transcripts while increasing markers of differentiation. (A)Western blot analysis of acid extracted histones from BB-p53nOPCs treated with EZH21 or DMSO for 48 hours and probed with antibodies specific for H3K27me3 and total H3. (B) Boxplot represents the ratio of H3K27me3 over total H3 calculated by dividing the intensity of the H3K27me3 bands by the intensity of total H3 bands from panel 4A data. Margins represent the lower and higher quartiles and ends minimum and maximum values of the data. DMSO BB-p53n mean normalized H3K27me3 = 1.17 +/-0.11 SEM. EZH2i BB-p53n mean normalized H3K27me3 = 0.67 +/-0.05 SEM. Significance calculated by one-tail student t-test (*p-*value = 0.017). (C) Heatmap of gene expression data of selected differentially expressed genes comparing p53n in blue columns, vehicle treated (DMSO) BB-p53n in orange columns, and EZH2 inhibitor treated BB-p53n OPCs in purple columns. All samples done in triplicate. Genes represented in rows and high relative gene expression is red while low relative gene expression is blue. Clusters identified by k-means implemented on the full transcriptome. Z-scores calculated with normalized tpm values. (D) Dotplots of overrepresentation analysis on genes from the three clusters labelled in panel C. Analysis run against the gene ontology database. Gene ratio on the x-axis represents the number of genes matching between input gene list and ontology gene set divided by total number of genes in input list. Blue to red color gradient per dot represents *adjusted p*-value of overrepresentation analysis, blue is higher and red is lower *p-*value. Size of dots represents number of genes matching between input gene list and ontology gene set. (E) Bar graph of RT-qPCR data targeting cell cycle control genes, Atad2 and Ccne2. Y-axis represents log_2_ fold change of ΔΔ*Ct* values normalized to housekeeping genes done in triplicate. Significance calculated by one-way ANOVA. * = *p-*value < 0.05, ** = *p-*value < 0.005. (F) Bar graph of RT-qPCR data targeting OPC differentiation control genes, Prmt5 and Erbb2. Y-axis represents log_2_ fold change of ΔΔ*Ct* values normalized to housekeeping genes done in triplicate. Significance calculated by one-way ANOVA. * = *p-*value < 0.05, ** = *p-*value < 0.005.

**FIGURE 5:**
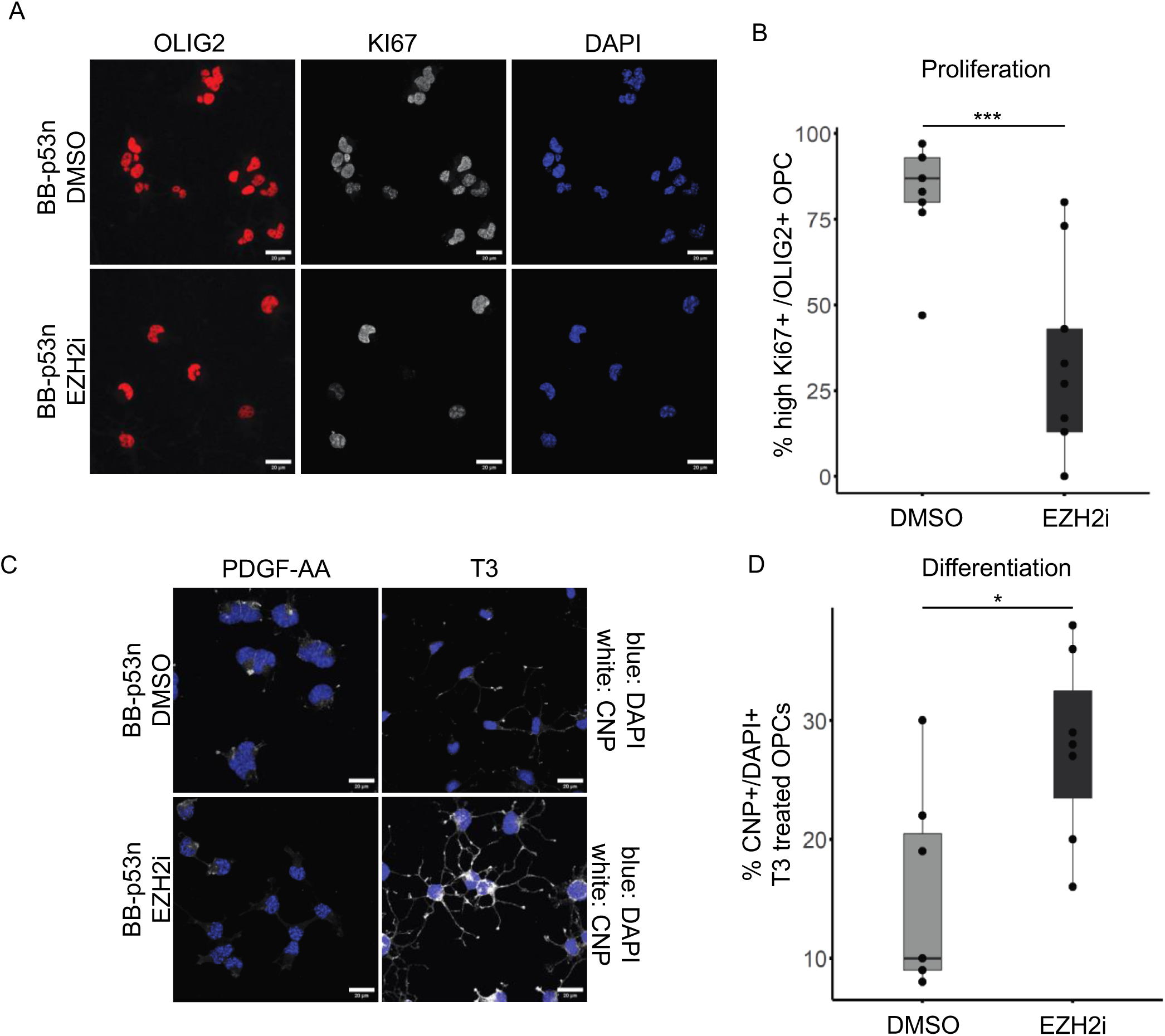
Reduction of H3K27me3 in BB-p53null OPCs by pharmacological inhibition of EZH2, reduces their proliferation and increases their differentiation ability. (A) Confocal images of DMSO vehicle treated BB-p53n and EZH2i treated BB-p53n OPCs. Cells treated for 48 hours and stained for OLIG2 (red), Ki67 (gray), and DAPI as a nuclear counterstain (blue). Scalebar = 20 μm. (B) Boxplot representing the percentage of proliferating (Ki67+/OLIG2+) cells described in panel 5A. Margins represent the lower and higher quartiles and ends minimum and maximum values of the data obtained from 9 independent determinations (n=30 cells counted for each experiment performed in triplicate in three independent biological samples). High Ki67 determined by nuclear mean fluorescent value above the median value of all samples. BB-p53n DMSO mean high Ki67 = 83% +/-4.99% SEM. BB-p53n EZH2i mean high Ki67 = 32% +/- 9.72% SEM. Significance two-way Student t-test (*p* value = *0.0006*). (C) Confocal images of 48 hours DMSO vehicle treated BB-p53n and 48 hours EZH2i treated BB-p53n OPCs cultured in 10 ng/mL PDGF-AA or 40ng/mL T3. After an additional 48 hours of PDGF-AA or T3 treatment, cells are fixed and stained for CNP (gray), and DAPI as a nuclear counterstain (blue). Scalebar = 20 μm. (D) Boxplot representing percent of CNP positive cells in T3 cultured BB-p53n OPCs. Margins represent the lower and higher quartiles and ends minimum and maximum values of the data obtained from 6 independent determinations (n=30 cells counted for each experiment performed in triplicate in two independent biological samples). CNP positive cells identified by setting the fluorescent threshold greater than the average of CNP fluorescence in PDGF-AA treated BB-p53n OPCs. BB-p53n DMSO mean %CNP+ = 15% +/-3.21% SEM. BB-p53n EZH2i mean %CNP+ = 28% +/-3.05% SEM. Significance two-tail Student t-test (*p* value = *0.016*)

Based on the transcriptional effects of EZH2i on BB-p53n OPCs, we asked whether also the functional properties of the cells were affected. We therefore assessed proliferation using immunocytochemistry and quantified the proportion of OLIG2+ cells that were also Ki67+. After 48 hour of EZH2i treatment, the BB-p53n OPCs showed significantly fewer Ki67+ cells (average percentage = 32 +/-9.72%) compared to the DMSO treated controls (average percentage= 83 +/-4.99%) (Fig5A,B). To further assess whether the same treatment also affected the ability of BB-p53n OPCs to differentiate, we removed mitogens and cultured cells in differentiation medium in the presence of either EZH2i or DMSO for an additional 48 hours. Differentiation was monitored using immunofluorescence with antibodies specific for the myelin protein for 2’,3’-Cyclic Nucleotide 3’ Phosphodiesterase (CNP)^51^, a marker for differentiated oligodendrocytes (Fig5C). The quantification of the CNP+ cells in EZH2i treated BB-p53n OPCs revealed an average of 28%+/-3.05 compared to the 15%+/-3.21% CNP+ cells in DMSO controls (Fig5D). Taken together, reduction of the H3K27me3 mark reduced the proliferation advantage induced by PDGF-BB in BB-p53n OPCs while also increasing their ability to differentiate when stimulated by T3.

## Discussion

This study was designed to address the early steps of gliomagenesis in oligodendrocyte progenitors, as these cells have been previously identified as the cell of origin of proneural gliomas^23,24^, a subtype of tumors characterized by the prevalence of OPC transcriptional signature^13,24^. Since specific genetic alterations have been identified for each subtype, with deletion of P53, NF1 or PTEN, IDH1 mutations and amplification of PDGF signaling reported in proneural gliomas, several groups adopted distinct strategies to create distinct animal models. Those included the use of the avian RCAS system^19,52^, the injection of retroviral vectors^22,53^, the combination of viral vector injection in transgenic mice^23,24,25^. Our results show that primary cultures of OPCs carrying specific genetic alterations, begin to show transcriptional and epigenetic changes *in vitro*, and will give rise to gliomas when injected into brain of recipient mice, providing a well-controlled experimental platform to test the effects of specific genetic alterations.

An open question that remains unanswered, however, is whether and how the initial genetic mutations occurring in OPC lead to transformation. This study addresses this question by investigating the effect of p53 deletion alone or in combination with PDGF-BB on the OPC transcriptome and histone post-translational modifications. We show here that OPCs carrying PDGF-BB overexpression and p53 deletion (BB-p53n OPC), are intrinsically distinct from OPCs carrying p53 deletion alone (p53n OPC) and are capable to induce the formation of gliomas 35 days after injection in the brain of recipient adult mice. Primary cultures of OPC carrying p53 deletion and PDGF overexpression were characterized by higher proliferative rates and decreased ability to differentiate than OPCs with p53 deletion alone. These findings are consistent with previous reports in mice with deletions of both *Trp53* and *Nf1* in adult OPC, where the premalignant phase of OPCs was characterized by increased proliferation and decreased differentiation^25^, thereby suggesting that proliferative advantage and inability to differentiate are common steps in gliomagenesis. In addition, to begin deciphering the molecular mechanisms underlying these early changes occurring in premalignant cells, we conducted an unbiased histone proteomics analysis, followed by validation using confocal imaging and western blot analysis of histone extracts. Collectively, these experiments detected higher levels of the repressive H3K27me3 histone mark in the BB-p53n OPCs compared to p53n OPCs, a finding which was also consistent with the higher expression levels of enzymes responsible for the deposition of this mark (e.g. EZH2), and lower transcript levels of enzymes responsible for erasing the repressive H3K27me3 histone mark. At a transcriptional level, BB-p53n OPCs were characterized by higher expression levels of genes encoding for positive regulation of the cell cycle and lower levels of transcripts related to oligodendrocyte differentiation.

These data are consistent with reports in adult gliomas, where higher levels of EZH2^54,55^ and higher levels of H3K27me3^56,57^ have been associated with worse prognosis. In our study, the reduction of H3K27me3 levels in OPCs with p53 deletion and PDGF-BB overexpression, by treatment with a well-recognized EZH2 inhibitor (Tazemetostat), significantly reduced the expression of genes regulating cell division, while increasing the expression of genes related to differentiation, thereby providing a molecular rationale to previous studies on the prognostic value of H3K27me3^56,57^. Our data are also consistent with the beneficial effect of reduced EZH2 activity on decreasing proliferation and reducing tumor growth in animal models^58,59,60^ and decreasing proliferation and migration in adult human glioma cells^61^.

Our results, however, differ from those reported for pediatric diffuse pontine intrinsic gliomas, characterized by the H3K27M mutation, where lysine residue K27 in histone H3 is replaced by a methionine, thereby resulting in overall decreased H3K27me3 mark^62,63,64,65,66,67^. Overall these data suggest that the effect of EZH2 inhibition on gliomagenesis is cell-type and developmental stage specific, and suitable for early pre-malignant states of transformation. In agreement with the cell specificity of the effect of EZH2 inhibition, and previous reports on the importance of EZH2 in OPC cell fate^68,69^ it is important to note that short term inhibition was shown to be beneficial for glioma survival, while long-term depletion resulted in cell fate switch and tumor progression^70^.

These data also highlight the limitations of treating cells with a single inhibitor, as the epigenomic landscape of cells cannot be singly ascribed to specific histone modifications. Accordingly, the overall effect of Tazemetostat in reversing the transcriptional differences between the BB-p53n and the p53n OPCs was only partial and affected genes regulating proliferation, resulting in elimination of the proliferative advantage and favoring differentiation. The partial reversal of the BB-p53n OPC transcriptome also suggests the possible existence of additional epigenetic differences with p53n OPC, such as potential differences in DNA and RNA methylation or hydroxymethylation and microRNAs, which could not be captured by the histone proteomic analysis. Overall, this study highlights the complexity of the early stages of gliomagenesis in OPC, identifies the H3K27me3 mark as critical for these early events, thereby identifying EZH2 inhibition as a potential strategy for early stages of glioma progression such as may be seen in diffuse low-grade gliomas.

## Limitations of the study

A potential limitation of this study is that we did not characterize the entire extent of the epigenetic landscape, as potential differences in DNA methylation, lnc RNAs and microRNAs could also be affected by the genetic alterations and contribute to the intrinsic differences between p53n and BB-p53n OPCs.

## Acknowledgments.

This work was performed thanks to grants from the National Institute of Health to Patrizia Casaccia (R35 - NS111604), to Peter Canoll and Patrizia Casaccia (5R01NS052738) and to Benjamin Garcia (P01CA196539; R01HD106051).

## Author contributions

Patrizia Casaccia conceived the project with Peter Canoll. She coordinated and supervised the work, wrote the final version of the manuscript, together with DH. All the experiments in cultured OPCs were performed by DH, including the analysis of the RNA Seq datasets. The animal injections and *in vivo* studies were carried out by AM. The histone proteomic analysis was conducted by BG and NVB. The figures were prepared by DH. All the authors provide comments to the final draft of the manuscript.

## Inclusion and diversity

We worked to ensure sex balance selection in the selection of non-human subjects. While citing references scientifically relevant for this work, we also actively worked to promote gender balance in our reference list.

## STAR METHODS

### RESOURCE AVAILABILITY

#### Lead contact

Further information and requests for resources and reagents should be directed to and will be fulfilled by the lead contact, Patrizia Casaccia.

#### Materials availability

Mouse lines used in this study are subject to MTA from the original investigator.

#### Data and code availability

RNA-Seq data have been deposited at NCBI GEO Repository and are publicly available as of the date of publication. Accession numbers are listed in the key resources table. All data reported in this study and any additional information required to reanalyze the data reported in this study are available from the lead contact upon request.

### EXPERIMENTAL MODEL AND SUBJECT DETAILS

#### OPC isolation, culture, infection

Primary mouse OPCs were isolated from the brain of *Trp53fl/fl* C57BL/6 (JAX:008462) mice at postnatal day 5-7 by with a rat anti-mouse CD140a antibody (Fisher Cat# 558774), recognizing PDGFRα, as previously described^68^ and were cultured in SATO medium (Dulbecco’s modified Eagle’s medium, 10 mg/ml bovine serum albumin (BSA), 10 mg/ml apotransferrin, 1.6 mg/ml putrescine, 6 ng/ml progesterone, 4 μg/ml selenium, 5 mg/ml insulin, 1 mM sodium pyruvate, 2 mM l-glutamine, 100 U/ml penicillin, 100 g/ml streptomycin, 5 mg/ml N-acetyl-cysteine, Trace Element B, 10 μg/ml biotin, 50 mM forskolin) supplemented with PDGF-AA (10 ng/ml) and basic fibroblast growth factor (bFGF) (20 ng/ml). *Trp53fl/fl* OPCs were then infected with an X-IRES-CRE or PDGFB-IRES-CRE retrovirus to obtain *Trp53−/−* (p53n) and *Trp53-/-* PDGFB expressing (BB-p53n) OPCs. Retrovirus production is described in Lei, 2011^24^. In brief, plasmid expressing VSVG and viral plasmid PDGFB-IRES-CRE were mixed with CaCl_2._ 2X HBS (Hepes Buffered Saline) was added to the mix, which was overlaid onto the 293GP cells. After 48 hours, the virus enriched media is filtered and concentrated. 500,000 primary mouse OPCs were plated into a 35mm plate before virus infection. Virus infection was performed by adding the concentrated virus directly into the tissue culture medium. The virus-containing medium was replaced by fresh media 12h after infection. Infected cells were harvested 6 days after infection for experimental analysis. The experiment was independently replicated by two investigators in the lab.

#### OPC Tazemetostat treatment and Differentiation

OPCs are cultured on 10cm Poly-D-Lysine (PDL) treated dishes in SATO growth media. Tazemetostat (EPZ-6438 Selleck cat#S7128) is dissolved in DMSO to a stock concentration of 50mM. Treatment begins when drug is added to the media to a dilution to 5uM and vehicle controls are given DMSO alone. Tazemetostat and DMSO added media is replaced every 24 hours for a total of 48 hours of treatment. In differentiation assays, OPCs are first treated with Tazemetostat or DMSO for 48 hours in 10cm dishes before being replated on PDL coated coverslips (Electron Microscopy Sciences Cat#72294-12) at a concentration of 20,000 cells per coverslip. OPCs are kept in Tazemetostat or DMSO while on coverslips and stimulated with Triiodothyronine (T3 60 nM) (Sigma cat#T5516-1MG) to induce differentiation for an addition 48 hours or PDGF-AA (10ng/mL) and bFGF (20ng/mL) as controls.

#### Mutant OPC in vivo injections

Cell implantation was performed by stereotactic intracranial injection of 50,000 OPCs in 1 μL of SATO, at a flow rate of 0.25 μL/min with a Hamilton syringe, as described previously described^71^. C57BL/6J Mice (JAX:000664) were anesthetized with Ketamine/Xylazine (100 mg/kg and 10 mg/kg, respectively) and assessed for lack of reflexes by toe pinch. A burr hole was made with a 17-gauge needle 2 mm lateral and 2 mm anterior to the bregma.

#### Immunocytochemistry

List of antibodies and buffers used for immunocytochemistry and immunoblotting are provided in Key Resources Table. Cells for immunocytochemistry were seeded in 8-well chamber slides (Thermo Fisher Scientific, 154941PK) or on PDL coated coverslips (Electron Microscopy Sciences Cat#72294-12) and fixed with 4% paraformaldehyde (PFA) for 15 min at room temperature. Membranes were permeabilized with 0.1% (vol/vol) Triton X-100/PBS (Fisher Scientific cat#AAA16046AP). Incubation with blocking solution (PGBA 5% normal goat serum Vector Laboratories cat#S-1000-20) was performed at room temperature for 60 min. Primary antibodies were applied overnight at 4 °C or 1hr at room temperature followed by incubation of appropriate secondary antibodies conjugated with fluorophores. DAPI (life technologies Cat#D21490) in PBS is applied for 10 minutes and washed before mounting. Confocal images were captured using the Zeiss LSM-800 fluorescent microscope and Zen Blue software. Blinded quantification of the immunofluorescent intensity was done using Fiji/ImageJ^72^. Boxplots generated in R with ggplot2 package^73^.

#### Western Blotting

In vitro cells, cultured to 90% confluency, were lysed for protein with RIPA buffer (Thermofisher cat# 89900) supplemented with phosphatase (Sigma cat#P0044-5ML) and protease (Thermofisher cat#A32963) inhibitors. Histone extraction is described below. Concentrations measured by Biorad’s DC protein assay (Biorad cat# 5000112) and 20ug of total protein or 1ug of acid extracted histones were loaded into the well of SurePAGE 4-20% gradient gels (GenScript Cat#M00656) and run in accompanying running buffer (GenScript Cat#M00138). Transfers were done with PVDF membrane and TG buffer 20% methanol. Membranes were blocked and probed for targets before imaging on the Biorad chemidoc imaging system. Quantifications of western blot signals done on Fiji/ImageJ^72^. Boxplots generated in R with ggplot2 package^73^.

#### RNA Isolation

Each RNA sample isolated from a 90% confluent 10cm culture dish. In vitro cells are homogenized in 1mL of TRIZOL reagent (Fisher cat# 15596026) and incubated at room temperature for 5 minutes. Phase separation is initiated by 0.2mL chloroform: isoamyl alcohol (Invitrogen cat#15593-049) and disrupted by vortex and shaking. After centrifuging at 4C for 15 minutes at 12,000xg, the aqueous phase is removed and mixed with 0.5mL isopropyl alcohol. Samples are incubated for 10 minutes at room temperature for the RNA to precipitate and then centrifuged at 12,000xg for 10 minutes at 4C. The collected RNA pellet is washed with 75% cold ethanol before being centrifuged again and reconstituted in RNase free water. RNA quantification is initially done with Nanodrop One (ThermoScientific cat#ND-ONE-W). RNA used for sequencing is cleaned up with the RNeasy MinElute Cleanup Kit (Qiagen cat#74204).

#### RNA sequencing library preparation and read processing

100ng of total RNA was used for library prep using Kapa mRNA HyperPrep Kit (Roche cat#KK8580) and sequenced on a Novaseq 6000 sequencer configured to paired end 150. Raw reads were trimmed by trimmomatic^74^, arguments set to ILLUMINACLIP:Illumina-adapt.fa:2:30:10, SLIDINGWINDOW:4:15, MINLEN:50. Trimmed reads aligned to mm39 reference genome^75^ with the subjunc tool of the subread package^76^, arguments set to defaults. 27-46 million paired reads met quality checks and were used to count features with the featuresCounts tool of the subread package, annotation by Ensembl^77^ (Mus_musculus.GRCm39.108) with feature type set to exon.

#### RNA sequencing Analysis

Gene count matrices were imported to R for all downstream analysis. Raw counts were normalized to TPM values and converted to scaled z-scores to plot heatmaps using the pheatmap (v1.0.12) package^78^. Differential expression analysis was done with limma in the edgeR (v4.0.16) package^79^. GSEA was performed with the clusterProfiler^80,81^ (v4.10.0) package and Org.Mm.eg.db^82^ (v3.18.0) annotation database. Gene lists for GSEA^41^ were filtered by adjusted *p*-value less than 0.01, calculated by limma differential expression analysis. Curated G1/S and G2/M genesets obtained from Tirosh et al^42^ and other gene sets obtained from the Gene Ontology database^43,44^. All enrichment plots are generated with the enrichplot^83^ (v1.22.0) package. All volcano plots generated by ggplot2^73^. R code is available at (https://github.com/dhuang-ASRC/BBp53n_Submission)

#### RT-qPCR

cDNA is synthesized with 500ng of RNA, 4uL of qScript XLT cDNA Supermix (Quantabio cat# 95161-500), and RNase free water to a volume of 20uL per sample. Thermocycler is set according to qScript instructions. Each PCR reaction contains 5uL of cDNA (1ng/uL), 6uL Perfecta Sybr Green FastMix (Quantbio cat# 95072-012), 0.5uL of RNase free water, 0.5uL 10uM Primer mix (primer sequences in supplementary table 7). Each reaction was pipetted into a 385 well plate and run on the QuantStudio 7 Flex with Perfecta Sybr Green FastMix thermocycling instructions. Log_2_ fold change was calculated by Δ ΔCt method normalized first to the average Ct values of three house keeping genes (*Ppia, Rpl13a, Pgk1*) and then to p53n control values.

#### Histone extraction and LC-MS/MS analysis

BB-p53n and p53n OPCs were cultured in mitogen supplemented SATO media to a 90% confluency on 15cm dishes, The histones were extracted and prepared for chemical derivatization and digestion as described previously^46,47^. In brief, cells were lysed in nuclear isolation buffer supplemented with protease and histone deacetylase inhibitors. Histones were precipitated with 25% trichloroacetic acid overnight and washed with acetic acid before reconstituted with RNase free water. The lysine residues from histones were derivatized with the propionylation reagent (1:2 reagent:sample ratio) containing acetonitrile and propionic anhydride (3:1), and the solution pH was adjusted to 8.0 using ammonium hydroxide. The propionylation was performed twice and the samples were dried on speed vac. The derivatized histones were then digested with trypsin at a 1:50 ratio (wt/wt) in 50 mM ammonium bicarbonate buffer at room temperature overnight. The N-termini of histone peptides were derivatized with the propionylation reagent twice and dried on speed vac. The peptides were desalted with the self-packed C18 stage tip. The purified peptides were then dried and reconstituted in 0.1% formic acid. Histone peptides were separated with Vanquish Neo UHPLC system fitted with 75 μm i.d. x 15 cm fused silica columns (Polymicro Tech) packed with ReproSil-Pur 120 C18-AQ (3 μm, Dr. Maisch GmbH) and connected in line with a mass spectrometer (Thermo QE). The chromatography conditions generally consisted of a linear gradient from 2 to 45% solvent B (0.1% formic acid in 80% acetonitrile) in solvent A (0.1% formic acid in water) over 55 mins and then 45 to 99% solvent B over 5 mins at a flow rate of 300 nL/min. The mass spectrometer was programmed for data-independent acquisition (DIA). One acquisition cycle consisted of a full MS scan, 35 DIA MS/MS scans of 24 m/z isolation width starting from 295 m/z to reach 1100 m/z. Typically, full MS scans were acquired in the Orbitrap mass analyzer across 295–1100 m/z at a resolution of 70,000 in positive profile mode with a maximum injection time of 50 ms and an AGC target of 1e6. MS/MS data from HCD fragmentation was collected in the ion trap (when available) or the Orbitrap. These scans typically used an NCE of 30, an AGC target of 2e5, and a maximum injection time of 60 ms. Histone MS data were analyzed with EpiProfile^45^.

#### Histone ratios analysis

Ratio values were converted to scaled z-scores for selected histone modifications in R and heatmaps were plotted in R with the ComplexHeatmap^84,85^ (v2.18.0) package.

## REFERENCES

1. Wang, Q., Hu, B., Hu, X., Kim, H., Squatrito, M., Scarpace, L., deCarvalho, A. C., Lyu, S., Li, P., Li, Y., Barthel, F., Cho, H. J., Lin, Y.-H., Satani, N., Martinez-Ledesma, E., Zheng, S., Chang, E., Sauvé, C.-E. G., Olar, A.,… Verhaak, R. G. W. (2017). Tumor Evolution of Glioma-Intrinsic Gene Expression Subtypes Associates with Immunological Changes in the Microenvironment. Cancer Cell, 32(1), 42–56.e6. 10.1016/j.ccell.2017.06.003

2. Patel, A. P., Tirosh, I., Trombetta, J. J., Shalek, A. K., Gillespie, S. M., Wakimoto, H., Cahill, D. P., Nahed, B. V., Curry, W. T., Martuza, R. L., Louis, D. N., Rozenblatt-Rosen, O., Suvà, M. L., Regev, A., & Bernstein, B. E. (2014). Single-cell RNA-seq highlights intratumoral heterogeneity in primary glioblastoma. Science, 344(6190), 1396–1401. 10.1126/science.1254257

3. Parker, N. R., Khong, P., Parkinson, J. F., Howell, V. M., & Wheeler, H. R. (2015). Molecular Heterogeneity in Glioblastoma: Potential Clinical Implications. Frontiers in Oncology, 5. https://www.frontiersin.org/articles/10.3389/fonc.2015.00055

4. Yuan, J., Levitin, H. M., Frattini, V., Bush, E. C., Boyett, D. M., Samanamud, J., Ceccarelli, M., Dovas, A., Zanazzi, G., Canoll, P., Bruce, J. N., Lasorella, A., Iavarone, A., & Sims, P. A. (2018). Single-cell transcriptome analysis of lineage diversity in high-grade glioma. Genome Medicine, 10(1), 57. 10.1186/s13073-018-0567-9.

5. Al-Dalahmah, O., Argenziano, M. G., Kannan, A., Mahajan, A., Furnari, J., Paryani, F., Boyett, D., Save, A., Humala, N., Khan, F., Li, J., Lu, H., Sun, Y., Tuddenham, J. F., Goldberg, A. R., Dovas, A., Banu, M. A., Sudhakar, T., Bush, E.,… Canoll, P. (2023). Re-convolving the compositional landscape of primary and recurrent glioblastoma reveals prognostic and targetable tissue states. Nature Communications, 14(1), 2586. 10.1038/s41467-023-38186-1

6. Klughammer, J., Kiesel, B., Roetzer, T., Fortelny, N., Kuchler, A., Nenning, K.-H., Furtner, J., Sheffield, N. C., Datlinger, P., Peter, N., Nowosielski, M., Augustin, M., Mischkulnig, M., Ströbel, T., Alpar, D., Ergüner, B., Senekowitsch, M., Moser, P., Freyschlag, C. F.,… Bock, C. (2018). The DNA methylation landscape of glioblastoma disease progression shows extensive heterogeneity in time and space. Nature Medicine, 24(10), 1611–1624. 10.1038/s41591-018-0156-x

7. Mack, S. C., Singh, I., Wang, X., Hirsch, R., Wu, Q., Villagomez, R., Bernatchez, J. A., Zhu, Z., Gimple, R. C., Kim, L. J. Y., Morton, A., Lai, S., Qiu, Z., Prager, B. C., Bertrand, K. C., Mah, C., Zhou, W., Lee, C., Barnett, G. H.,… Rich, J. N. (2019). Chromatin landscapes reveal developmentally encoded transcriptional states that define human glioblastoma. The Journal of Experimental Medicine, 216(5), 1071–1090. 10.1084/jem.20190196

8. Chinot, O. L., Wick, W., Mason, W., Henriksson, R., Saran, F., Nishikawa, R., Carpentier, A. F., Hoang-Xuan, K., Kavan, P., Cernea, D., Brandes, A. A., Hilton, M., Abrey, L., & Cloughesy, T. (2014). Bevacizumab plus radiotherapy-temozolomide for newly diagnosed glioblastoma. The New England Journal of Medicine, 370(8), 709–722. 10.1056/NEJMoa1308345

9. Gilbert, M. R., Dignam, J. J., Armstrong, T. S., Wefel, J. S., Blumenthal, D. T., Vogelbaum, M. A., Colman, H., Chakravarti, A., Pugh, S., Won, M., Jeraj, R., Brown, P. D., Jaeckle, K. A., Schiff, D., Stieber, V. W., Brachman, D. G., Werner-Wasik, M., Tremont-Lukats, I. W., Sulman, E. P.,… Mehta, M. P. (2014). A Randomized Trial of Bevacizumab for Newly Diagnosed Glioblastoma. New England Journal of Medicine, 370(8), 699–708. 10.1056/NEJMoa1308573

10. Stupp, R., Mason, W. P., van den Bent, M. J., Weller, M., Fisher, B., Taphoorn, M. J. B., Belanger, K., Brandes, A. A., Marosi, C., Bogdahn, U., Curschmann, J., Janzer, R. C., Ludwin, S. K., Gorlia, T., Allgeier, A., Lacombe, D., Cairncross, J. G., Eisenhauer, E., & Mirimanoff, R. O. (2005). Radiotherapy plus Concomitant and Adjuvant Temozolomide for Glioblastoma. New England Journal of Medicine, 352(10), 987–996. 10.1056/NEJMoa043330

11. Nigro, J. M., Misra, A., Zhang, L., Smirnov, I., Colman, H., Griffin, C., Ozburn, N., Chen, M., Pan, E., Koul, D., Yung, W. K. A., Feuerstein, B. G., & Aldape, K. D. (2005). Integrated array-comparative genomic hybridization and expression array profiles identify clinically relevant molecular subtypes of glioblastoma. Cancer Research, 65(5), 1678–1686. 10.1158/0008-5472.CAN-04-2921

12. Liang, Y., Diehn, M., Watson, N., Bollen, A. W., Aldape, K. D., Nicholas, M. K., Lamborn, K. R., Berger, M. S., Botstein, D., Brown, P. O., & Israel, M. A. (2005). Gene expression profiling reveals molecularly and clinically distinct subtypes of glioblastoma multiforme. Proceedings of the National Academy of Sciences, 102(16), 5814–5819. 10.1073/pnas.0402870102

13. Verhaak, R. G. W., Hoadley, K. A., Purdom, E., Wang, V., Qi, Y., Wilkerson, M. D., Miller, C. R., Ding, L., Golub, T., Mesirov, J. P., Alexe, G., Lawrence, M., O’Kelly, M., Tamayo, P., Weir, B. A., Gabrie, S., Winckler, W., Gupta, S., Jakkula, L.,… Hayes, D. N. (2010). An integrated genomic analysis identifies clinically relevant subtypes of glioblastoma characterized by abnormalities in PDGFRA, IDH1, EGFR and NF1. Cancer Cell, 17(1), 98. 10.1016/j.ccr.2009.12.020

14. Gensert, J. M., & Goldman, J. E. (2001). Heterogeneity of cycling glial progenitors in the adult mammalian cortex and white matter. Journal of Neurobiology, 48(2), 75–86.

15. Bergles, D. E., & Richardson, W. D. (2016). Oligodendrocyte Development and Plasticity. Cold Spring Harbor Perspectives in Biology, 8(2), a020453. 10.1101/cshperspect.a020453

16. Nishiyama, A., Suzuki, R., & Zhu, X. (2014). NG2 cells (polydendrocytes) in brain physiology and repair. Frontiers in Neuroscience, 0. 10.3389/fnins.2014.00133

17. The Cancer Genome Atlas Research Network. Comprehensive genomic characterization defines human glioblastoma genes and core pathways. Nature 455, 1061–1068 (2008). 10.1038/nature07385

18. Parsons, D. W., Jones, S., Zhang, X., Lin, J. C.-H., Leary, R. J., Angenendt, P., Mankoo, P., Carter, H., Siu, I.-M., Gallia, G. L., Olivi, A., McLendon, R., Rasheed, B. A., Keir, S., Nikolskaya, T., Nikolsky, Y., Busam, D. A., Tekleab, H., Diaz, L. A.,… Kinzler, K. W. (2008). An integrated genomic analysis of human glioblastoma multiforme. Science (New York, N.Y.), 321(5897), 1807–1812. 10.1126/science.1164382

19. Dai, C., Celestino, J. C., Okada, Y., Louis, D. N., Fuller, G. N., & Holland, E. C. (2001). PDGF autocrine stimulation dedifferentiates cultured astrocytes and induces oligodendrogliomas and oligoastrocytomas from neural progenitors and astrocytes in vivo. Genes & Development, 15(15), 1913–1925. 10.1101/gad.903001

20. Hambardzumyan, D., Amankulor, N. M., Helmy, K. Y., Becher, O. J., & Holland, E. C. (2009). Modeling Adult Gliomas Using RCAS/t-va Technology. Translational Oncology, 2(2), 89–95. https://www.ncbi.nlm.nih.gov/pmc/articles/PMC2670576/

21. Lindberg, N., Kastemar, M., Olofsson, T., Smits, A., & Uhrbom, L. (2009). Oligodendrocyte progenitor cells can act as cell of origin for experimental glioma. Oncogene, 28(23), 2266–2275. 10.1038/onc.2009.76

22. Assanah, M., Lochhead, R., Ogden, A., Bruce, J., Goldman, J., & Canoll, P. (2006). Glial Progenitors in Adult White Matter Are Driven to Form Malignant Gliomas by Platelet-Derived Growth Factor-Expressing Retroviruses. Journal of Neuroscience, 26(25), 6781–6790. 10.1523/JNEUROSCI.0514-06.2006

23. Liu, C., Sage, J. C., Miller, M. R., Verhaak, R. G. W., Hippenmeyer, S., Vogel, H., Foreman, O., Bronson, R. T., Nishiyama, A., Luo, L., & Zong, H. (2011). Mosaic Analysis with Double Markers Reveals Tumor Cell of Origin in Glioma. Cell, 146(2), 209–221. 10.1016/j.cell.2011.06.014

24. Lei, L., Sonabend, A. M., Guarnieri, P., Soderquist, C., Ludwig, T., Rosenfeld, S., Bruce, J. N., & Canoll, P. (2011). Glioblastoma Models Reveal the Connection between Adult Glial Progenitors and the Proneural Phenotype. PLOS ONE, 6(5), e20041. 10.1371/journal.pone.0020041

25. Galvao, R. P., Kasina, A., McNeill, R. S., Harbin, J. E., Foreman, O., Verhaak, R. G. W., Nishiyama, A., Miller, C. R., & Zong, H. (2014). Transformation of quiescent adult oligodendrocyte precursor cells into malignant glioma through a multistep reactivation process. Proceedings of the National Academy of Sciences, 111(40), E4214–E4223. 10.1073/pnas.1414389111

26. Noble, M., Murray, K., Stroobant, P., Waterfield, M. D., & Riddle, P. (1988). Platelet-derived growth factor promotes division and motility and inhibits premature differentiation of the oligodendrocyte/type −2 astrocyte progenitor ceil. Nature, 333(6173), 560–562. 10.1038/333560a0

27. Raff, M. C., Lillien, L. E., Richardson, W. D., Burne, J. F., & Noble, M. D. (1988). Platelet-derived growth factor from astrocytes drives the clock that times oligodendrocyte development in culture. Nature, 333(6173), 562–565. 10.1038/333562a0

28. Pringle, N., Collarini, E. J., Mosley, M. J., Heldin, C. H., Westermark, B., & Richardson, W. D. (1989). PDGF A chain homodimers drive proliferation of bipotential (O-2A) glial progenitor cells in the developing rat optic nerve. The EMBO Journal, 8(4), 1049–1056. 10.1002/j.1460-2075.1989.tb03472.x

29. Guo, P., Hu, B., Gu, W., Xu, L., Wang, D., Huang, H.-J. S., Cavenee, W. K., & Cheng, S.-Y. (2003). Platelet-Derived Growth Factor-B Enhances Glioma Angiogenesis by Stimulating Vascular Endothelial Growth Factor Expression in Tumor Endothelia and by Promoting Pericyte Recruitment. The American Journal of Pathology, 162(4), 1083–1093. 10.1016/S0002-9440(10)63905-3

30. Zheng, Y., Yamamoto, S., Ishii, Y., Sang, Y., Hamashima, T., Van De, N., Nishizono, H., Inoue, R., Mori, H., & Sasahara, M. (2016). Glioma-Derived Platelet-Derived Growth Factor-BB Recruits Oligodendrocyte Progenitor Cells via Platelet-Derived Growth Factor Receptor-α and Remodels Cancer Stroma. The American Journal of Pathology, 186(5), 1081–1091. 10.1016/j.ajpath.2015.12.020

31. Betsholtz, C., Johnsson, A., Heldin, C.-H., Westermark, B., Lind, P., Urdea, M. S., Eddy, R., Shows, T. B., Philpott, K., Mellor, A. L., Knott, T. J., & Scott, J. (1986). cDNA sequence and chromosomal localization of human platelet-derived growth factor A-chain and its expression in tumour cell lines. Nature, 320(6064), 695–699. 10.1038/320695a0

32. Nistér, M., Claesson-Welsh, L., Eriksson, A., Heldin, C. H., & Westermark, B. (1991). Differential expression of platelet-derived growth factor receptors in human malignant glioma cell lines. The Journal of Biological Chemistry, 266(25), 16755–16763.

33. Nistér, M., Heldin, C.-H., & Westermark, B. (1986). Clonal Variation in the Production of a Platelet-derived Growth Factor-like Protein and Expression of Corresponding Receptors in a Human Malignant Glioma1. Cancer Research, 46(1), 332–340.

34. Lokker, N. A., Sullivan, C. M., Hollenbach, S. J., Israel, M. A., & Giese, N. A. (2002). Platelet-derived growth factor (PDGF) autocrine signaling regulates survival and mitogenic pathways in glioblastoma cells: evidence that the novel PDGF-C and PDGF-D ligands may play a role in the development of brain tumors. Cancer Research, 62(13), 3729–3735.

35. Kim, Y., Kim, E., Wu, Q., Guryanova, O., Hitomi, M., Lathia, J. D., Serwanski, D., Sloan, A. E., Weil, R. J., Lee, J., Nishiyama, A., Bao, S., Hjelmeland, A. B., & Rich, J. N. (2012). Platelet-derived growth factor receptors differentially inform intertumoral and intratumoral heterogeneity. Genes & Development, 26(11), 1247–1262. 10.1101/gad.193565.112

36. Thijssen, V. L., Paulis, Y. W., Nowak-Sliwinska, P., Deumelandt, K. L., Hosaka, K., Soetekouw, P. M., Cimpean, A. M., Raica, M., Pauwels, P., van den Oord, J. J., Tjan-Heijnen, V. C., Hendrix, M. J., Heldin, C.-H., Cao, Y., & Griffioen, A. W. (2018). Targeting PDGF-mediated recruitment of pericytes blocks vascular mimicry and tumor growth. The Journal of Pathology, 246(4), 447–458. 10.1002/path.5152

37. Marino, S., Vooijs, M., van Der Gulden, H., Jonkers, J., & Berns, A. (2000). Induction of medulloblastomas in p53-null mutant mice by somatic inactivation of Rb in the external granular layer cells of the cerebellum. Genes & Development, 14(8), 994–1004.

38. Sonabend, A. M., Yun, J., Lei, L., Leung, R., Soderquist, C., Crisman, C., Gill, B. J., Carminucci, A., Sisti, J., Castelli, M., Sims, P. A., Bruce, J. N., & Canoll, P. (2013). Murine Cell Line Model of Proneural Glioma for Evaluation of Anti-Tumor Therapies. Journal of Neuro-Oncology, 112(3), 375–382. 10.1007/s11060-013-1082-x

39. Zhou, Q., Wang, S., & Anderson, D. J. (2000). Identification of a novel family of oligodendrocyte lineage-specific basic helix-loop-helix transcription factors. Neuron, 25(2), 331–343. 10.1016/s0896-6273(00)80898-3

40. Miller, I., Min, M., Yang, C., Tian, C., Gookin, S., Carter, D., & Spence, S. L. (2018). Ki67 is a Graded Rather than a Binary Marker of Proliferation versus Quiescence. Cell Reports, 24(5), 1105–1112.e5. 10.1016/j.celrep.2018.06.110

41. Subramanian, A., Tamayo, P., Mootha, V. K., Mukherjee, S., Ebert, B. L., Gillette, M. A., Paulovich, A., Pomeroy, S. L., Golub, T. R., Lander, E. S., & Mesirov, J. P. (2005). Gene set enrichment analysis: A knowledge-based approach for interpreting genome-wide expression profiles. Proceedings of the National Academy of Sciences, 102(43), 15545–15550. 10.1073/pnas.0506580102

42. Tirosh, I., Venteicher, A. S., Hebert, C., Escalante, L. E., Patel, A. P., Yizhak, K., Fisher, J. M., Rodman, C., Mount, C., Filbin, M. G., Neftel, C., Desai, N., Nyman, J., Izar, B., Luo, C. C., Francis, J. M., Patel, A. A., Onozato, M. L., Riggi, N.,… Suvà, M. L. (2016). Single-cell RNA-seq supports a developmental hierarchy in human oligodendroglioma. Nature, 539(7628), 309–313. 10.1038/nature20123

43. Ashburner, M., Ball, C. A., Blake, J. A., Botstein, D., Butler, H., Cherry, J. M., Davis, A. P., Dolinski, K., Dwight, S. S., Eppig, J. T., Harris, M. A., Hill, D. P., Issel-Tarver, L., Kasarskis, A., Lewis, S., Matese, J. C., Richardson, J. E., Ringwald, M., Rubin, G. M., & Sherlock, G. (2000). Gene Ontology: tool for the unification of biology. Nature Genetics, 25(1), 25–29. 10.1038/75556

44. The Gene Ontology Consortium, Aleksander, S. A., Balhoff, J., Carbon, S., Cherry, J. M., Drabkin, H. J., Ebert, D., Feuermann, M., Gaudet, P., Harris, N. L., Hill, D. P., Lee, R., Mi, H., Moxon, S., Mungall, C. J., Muruganugan, A., Mushayahama, T., Sternberg, P. W., Thomas, P. D.,… Westerfield, M. (2023). The Gene Ontology knowledgebase in 2023. Genetics, 224(1), iyad031. 10.1093/genetics/iyad031

45. Yuan, Z.-F., Lin, S., Molden, R. C., Cao, X.-J., Bhanu, N. V., Wang, X., Sidoli, S., Liu, S., & Garcia, B. A. (2015). EpiProfile Quantifies Histone Peptides With Modifications by Extracting Retention Time and Intensity in High-resolution Mass Spectra. Molecular & Cellular Proteomics: MCP, 14(6), 1696–1707. 10.1074/mcp.M114.046011

46. Bhanu, N. V., Sidoli, S., & Garcia, B. A. (2020). A Workflow for Ultra-rapid Analysis of Histone Post-translational Modifications with Direct-injection Mass Spectrometry. Bio-Protocol, 10(18), e3756. 10.21769/BioProtoc.3756

47. Sidoli, S., Bhanu, N. V., Karch, K. R., Wang, X., & Garcia, B. A. (2016). Complete Workflow for Analysis of Histone Post-translational Modifications Using Bottom-up Mass Spectrometry: From Histone Extraction to Data Analysis. Journal of Visualized Experiments: JoVE, 111, 54112. 10.3791/54112

48. Koo, S. J., Fernández-Montalván, A. E., Badock, V., Ott, C. J., Holton, S. J., von Ahsen, O., Toedling, J., Vittori, S., Bradner, J. E., & Gorjánácz, M. (2016). ATAD2 is an epigenetic reader of newly synthesized histone marks during DNA replication. Oncotarget, 7(43), 70323–70335. 10.18632/oncotarget.11855

49. Scaglione, A., Patzig, J., Liang, J., Frawley, R., Bok, J., Mela, A., Yattah, C., Zhang, J., Teo, S. X., Zhou, T., Chen, S., Bernstein, E., Canoll, P., Guccione, E., & Casaccia, P. (2018). PRMT5-mediated regulation of developmental myelination. Nature Communications, 9, 2840. 10.1038/s41467-018-04863-9

50. Park, S.-K., Miller, R., Krane, I., & Vartanian, T. (2001). The erbB2 gene is required for the development of terminally differentiated spinal cord oligodendrocytes. Journal of Cell Biology, 154(6), 1245–1258. 10.1083/jcb.200104025

51. Raasakka, A., & Kursula, P. (2014). The myelin membrane-associated enzyme 2′,3′-cyclic nucleotide 3′-phosphodiesterase: on a highway to structure and function. Neuroscience Bulletin, 30(6), 956–966. 10.1007/s12264-013-1437-5

52. Hesselager, G., Uhrbom, L., Westermark, B., & Nistér, M. (2003). Complementary Effects of Platelet-derived Growth Factor Autocrine Stimulation and p53 or Ink4a-Arf Deletion in a Mouse Glioma Model1. Cancer Research, 63(15), 4305–4309.

53. Shih, A. H., Dai, C., Hu, X., Rosenblum, M. K., Koutcher, J. A., & Holland, E. C. (2004). Dose-dependent effects of platelet-derived growth factor-B on glial tumorigenesis. Cancer Research, 64(14), 4783–4789. 10.1158/0008-5472.CAN-03-3831\

54. Zhang, Y., Yu, X., Chen, L., Zhang, Z., & Feng, S. (2016). EZH2 overexpression is associated with poor prognosis in patients with glioma. Oncotarget, 8(1), 565–573. 10.18632/oncotarget.13478

55. Chen, Y., Hou, S., Jiang, R., Sun, J., Cheng, C., & Qian, Z. (2021). EZH2 is a potential prognostic predictor of glioma. Journal of Cellular and Molecular Medicine, 25(2), 925 –936. 10.1111/jcmm.16149

56. Ammendola, S., Caldonazzi, N., Simbolo, M., Piredda, M. L., Brunelli, M., Poliani, P. L., Pinna, G., Sala, F., Ghimenton, C., Scarpa, A., & Barresi, V. (2021). H3K27me3 immunostaining is diagnostic and prognostic in diffuse gliomas with oligodendroglial or mixed oligoastrocytic morphology. Virchows Archiv, 479(5), 987–996. 10.1007/s00428-021-03134-1

57. Filipski, K., Braun, Y., Zinke, J., Roller, B., Baumgarten, P., Wagner, M., Senft, C., Zeiner, P. S., Ronellenfitsch, M. W., Steinbach, J. P., Plate, K. H., Gasparoni, G., Mittelbronn, M., Capper, D., & Harter, P. N. (2019). Lack of H3K27 trimethylation is associated with 1p/19q codeletion in diffuse gliomas. Acta Neuropathologica, 138(2), 331–334. 10.1007/s00401-019-02025-9

58. Sprinzen, L., Garcia, F., Mela, A., Lei, L., Upadhyayula, P., Mahajan, A., Humala, N., Manier, L., Caprioli, R., Quiñones-Hinojosa, A., Casaccia, P., & Canoll, P. (2024). EZH2 Inhibition Sensitizes IDH1R132H-Mutant Gliomas to Histone Deacetylase Inhibitor. Cells, 13(3), 219. 10.3390/cells13030219

59. Karami Fath, M., Babakhaniyan, K., Anjomrooz, M., Jalalifar, M., Alizadeh, S. D., Pourghasem, Z., Abbasi Oshagh, P., Azargoonjahromi, A., Almasi, F., Manzoor, H. Z., Khalesi, B., Pourzardosht, N., Khalili, S., & Payandeh, Z. (2022). Recent Advances in Glioma Cancer Treatment: Conventional and Epigenetic Realms. Vaccines, 10(9), 1448. 10.3390/vaccines10091448

60. Rugo, H. S., Jacobs, I., Sharma, S., Scappaticci, F., Paul, T. A., Jensen-Pergakes, K., & Malouf, G. G. (2020). The Promise for Histone Methyltransferase Inhibitors for Epigenetic Therapy in Clinical Oncology: A Narrative Review. Advances in Therapy, 37(7), 3059. 10.1007/s12325-020-01379-x

61. Del Moral-Morales, A., González-Orozco, J. C., Hernández-Vega, A. M., Hernández-Ortega, K., Peña-Gutiérrez, K. M., & Camacho-Arroyo, I. (2022). EZH2 Mediates Proliferation, Migration, and Invasion Promoted by Estradiol in Human Glioblastoma Cells. Frontiers in Endocrinology, 13. 10.3389/fendo.2022.703733

62. Bender, S., Tang, Y., Lindroth, A. M., Hovestadt, V., Jones, D. T. W., Kool, M., Zapatka, M., Northcott, P. A., Sturm, D., Wang, W., Radlwimmer, B., Højfeldt, J. W., Truffaux, N., Castel, D., Schubert, S., Ryzhova, M., Şeker-Cin, H., Gronych, J., Johann, P. D.,… Pfister, S. M. (2013). Reduced H3K27me3 and DNA Hypomethylation Are Major Drivers of Gene Expression in K27M Mutant Pediatric High-Grade Gliomas. Cancer Cell, 24(5), 660–672. 10.1016/j.ccr.2013.10.006

63. Dhar, S., Gadd, S., Patel, P., Vaynshteyn, J., Raju, G. P., Hashizume, R., Brat, D. J., & Becher, O. J. (2022). A tumor suppressor role for EZH2 in diffuse midline glioma pathogenesis. Acta Neuropathologica Communications, 10(1), 47. 10.1186/s40478-022-01336-5

64. Harutyunyan, A. S., Chen, H., Lu, T., Horth, C., Nikbakht, H., Krug, B., Russo, C., Bareke, E., Marchione, D. M., Coradin, M., Garcia, B. A., Jabado, N., & Majewski, J. (2020). H3K27M in Gliomas Causes a One-Step Decrease in H3K27 Methylation and Reduced Spreading within the Constraints of H3K36 Methylation. Cell Reports, 33(7), 108390. 10.1016/j.celrep.2020.108390

65. Harutyunyan, A. S., Krug, B., Chen, H., Papillon-Cavanagh, S., Zeinieh, M., De Jay, N., Deshmukh, S., Chen, C. C. L., Belle, J., Mikael, L. G., Marchione, D. M., Li, R., Nikbakht, H., Hu, B., Cagnone, G., Cheung, W. A., Mohammadnia, A., Bechet, D., Faury, D.,… Majewski, J. (2019). H3K27M induces defective chromatin spread of PRC2-mediated repressive H3K27me2/me3 and is essential for glioma tumorigenesis. Nature Communications, 10(1), 1262. 10.1038/s41467-019-09140-x

66. Lewis, P. W., Müller, M. M., Koletsky, M. S., Cordero, F., Lin, S., Banaszynski, L. A., Garcia, B. A., Muir, T. W., Becher, O. J., & Allis, C. D. (2013). Inhibition of PRC2 Activity by a Gain-of-Function H3 Mutation Found in Pediatric Glioblastoma. Science, 340(6134), 857–861. 10.1126/science.1232245

67. Wu, G., Broniscer, A., McEachron, T. A., Lu, C., Paugh, B. S., Becksfort, J., Qu, C., Ding, L., Huether, R., Parker, M., Zhang, J., Gajjar, A., Dyer, M. A., Mullighan, C. G., Gilbertson, R. J., Mardis, E. R., Wilson, R. K., Downing, J. R., Ellison, D. W.,… St. Jude Children’s Research Hospital–Washington University Pediatric Cancer Genome Project. (2012). Somatic histone H3 alterations in pediatric diffuse intrinsic pontine gliomas and non-brainstem glioblastomas. Nature Genetics, 44(3), 251–253. 10.1038/ng.1102

68. Liu, J., Magri, L., Zhang, F., Marsh, N. O., Albrecht, S., Huynh, J. L., Kaur, J., Kuhlmann, T., Zhang, W., Slesinger, P. A., & Casaccia, P. (2015). Chromatin Landscape Defined by Repressive Histone Methylation during Oligodendrocyte Differentiation. The Journal of Neuroscience, 35(1), 352–365. 10.1523/JNEUROSCI.2606-14.2015

69. Sher, F., Rößler, R., Brouwer, N., Balasubramaniyan, V., Boddeke, E., & Copray, S. (2008). Differentiation of Neural Stem Cells into Oligodendrocytes: Involvement of the Polycomb Group Protein Ezh2. STEM CELLS, 26(11), 2875–2883. 10.1634/stemcells.2008-0121

70. de Vries, N. A., Hulsman, D., Akhtar, W., de Jong, J., Miles, D. C., Blom, M., van Tellingen, O., Jonkers, J., & van Lohuizen, M. (2015). Prolonged Ezh2 Depletion in Glioblastoma Causes a Robust Switch in Cell Fate Resulting in Tumor Progression. Cell Reports, 10(3), 383–397. 10.1016/j.celrep.2014.12.028

71. Upadhyayula PS, Higgins DM, Mela A et al. Dietary restriction of cysteine and methionine sensitizes gliomas to ferroptosis and induces alterations in energetic metabolism. Nat Commun. 2023 Mar 2;14(1):1187. doi: 10.1038/s41467-023-36630-w. PMID: 36864031; PMCID: PMC9981683

72. Schindelin, J., Arganda-Carreras, I., Frise, E., Kaynig, V., Longair, M., Pietzsch, T., Preibisch, S., Rueden, C., Saalfeld, S., Schmid, B., Tinevez, J.-Y., White, D. J., Hartenstein, V., Eliceiri, K., Tomancak, P., & Cardona, A. (2012). Fiji: an open-source platform for biological-image analysis. Nature Methods, 9(7), 676–682. 10.1038/nmeth.2019

73. Wickham H (2016). Ggplot­2: Elegant Graphics for Data Analysis. Springer-Verlag New York. ISBN 978-3-319-24277-4, https://ggplot2.tidyverse.org.

74. Bolger, A. M., Lohse, M., & Usadel, B. (2014). Trimmomatic: a flexible trimmer for Illumina sequence data. Bioinformatics, 30(15), 2114. 10.1093/bioinformatics/btu170

75. Mus Musculus mm39 Genome Assembly. NIH: National Library of Medicine. (https://www.ncbi.nlm.nih.gov/datasets/genome/GCF_000001635.27/)

76. Liao, Y., Smyth, G. K., & Shi, W. (2013). The Subread aligner: fast, accurate and scalable read mapping by seed-and-vote. Nucleic Acids Research, 41(10), e108. 10.1093/nar/gkt214

77. Martin, F. J., Amode, M. R., Aneja, A., Austine-Orimoloye, O., Azov, A. G., Barnes, I., Becker, A., Bennett, R., Berry, A., Bhai, J., Bhurji, S. K., Bignell, A., Boddu, S., Branco Lins, P. R., Brooks, L., Ramaraju, S. B., Charkhchi, M., Cockburn, A., Da Rin Fiorretto, L.,… Flicek, P. (2023). Ensembl 2023. Nucleic Acids Research, 51(D1), D933–D941. 10.1093/nar/gkac958

78. Kolde, R. Pheatmap: pretty heatmaps. R Package Version 1.0.10. https://CRAN.R-project.org/package=pheatmap (2012).

79. Robinson, M. D., McCarthy, D. J., & Smyth, G. K. (2010). edgeR: a Bioconductor package for differential expression analysis of digital gene expression data. Bioinformatics, 26(1), 139–140. 10.1093/bioinformatics/btp616

80. Wu, T., Hu, E., Xu, S., Chen, M., Guo, P., Dai, Z., Feng, T., Zhou, L., Tang, W., Zhan, L., Fu, X., Liu, S., Bo, X., & Yu, G. (2021). clusterProfiler 4.0: A universal enrichment tool for interpreting omics data. The Innovation, 2(3), 100141. 10.1016/j.xinn.2021.100141

81. Yu, G., Wang, L.-G., Han, Y., & He, Q.-Y. (2012). clusterProfiler: an R Package for Comparing Biological Themes Among Gene Clusters. OMICS: A Journal of Integrative Biology, 16(5), 284–287. 10.1089/omi.2011.0118

82. Carlson M (2019). org.Mm.eg.db: Genome wide annotation for Mouse. R package version 3.8.2.

83. Yu G (2023). enrichplot: Visualization of Functional Enrichment Result. doi:10.18129/B9.bioc.enrichplot, R package version 1.22.0, https://bioconductor.org/packages/enrichplot.

84. Gu, Z. (2022). Complex heatmap visualization. IMeta, 1(3), e43. 10.1002/imt2.43

85. Gu, Z., Eils, R., & Schlesner, M. (2016). Complex heatmaps reveal patterns and correlations in multidimensional genomic data. Bioinformatics, 32(18), 2847–2849. 10.1093/bioinformatics/btw313

